# Cell-cycle coupled expression minimizes random fluctuations in gene product levels

**DOI:** 10.1101/052159

**Authors:** Mohammad Soltani, Abhyudai Singh

**Author notes:** Corresponding Author: Abhyudai Singh,.

## Abstract

Expression of many genes varies as a cell transitions through different cell-cycle stages. How coupling between stochastic expression and cell cycle impacts cell-to-cell variability (noise) in the level of protein is not well understood. We analyze a model, where a stable protein is synthesized in random bursts, and the frequency with which bursts occur varies within the cell cycle. Formulas quantifying the extent of fluctuations in the protein copy number are derived and decomposed into components arising from the cell cycle and stochastic processes. The latter stochastic component represents contributions from bursty expression and errors incurred during partitioning of molecules between daughter cells. These formulas reveal an interesting trade-off: cell-cycle dependencies that amplify the noise contribution from bursty expression also attenuate the contribution from partitioning errors. We investigate existence of optimum strategies for coupling expression to the cell cycle that minimize the stochastic component. Intriguingly, results show that a zero production rate throughout the cell cycle, with expression only occurring just before cell division minimizes noise from bursty expression for a fixed mean protein level. In contrast, the optimal strategy in the case of partitioning errors is to make the protein just after cell division. We provide examples of regulatory proteins that are expressed only towards the end of cell cycle, and argue that such strategies enhance robustness of cell-cycle decisions to the intrinsic stochasticity of gene expression.

## 1. Introduction

Advances in experimental technologies over the last decade have provided important insights into gene expression at a single-molecule and single-cell resolution. An important (but not surprising) revelation is the stochastic expression of genes inside individual cells across different organisms [1–11]. In many cases, stochastic expression is characterized by random burst-like synthesis of gene products during transcription and translation. At the transcriptional level, promoters randomly switch to an active state, producing a burst of RNAs before becoming inactive [12–17]. At the translational level, a relatively unstable mRNA degrades after synthesizing a burst of protein molecules [18–21]. Bursty expression drives intercellular variability (noise) in gene product levels across isogenic cells, significantly impacting biological pathways and phenotypes [22–29].

Mathematical models have played a key role in predicting the impact of bursty expression on noise in the level of a given protein. However, these studies have primarily relied on models where synthesis rates are assumed to be constant and invariant of cell-cycle processes. While such an assumption is clearly violated for cell-cycle regulated genes, replication-associate changes in gene dosage can alter expression parameters genome wide [30–33]. It is not clear how such cell-cycle dependent expression affects the stochastic dynamics of protein levels in single cells. To systematically investigate this question, we formulate a model where a cell passes through multiple cell-cycle stages from birth to division. Cell cycle is coupled to bursty expression of a stable protein and the rate at which bursts occur depend arbitrarily on the cell-cycle stage (Fig. 1). In addition to stochastic expression in bursts, the model incorporates other physiological noise sources, such as, variability in the duration of cell-cycle times and random partitioning of molecules between daughter cells at the time of division [34–42].

**Figure 1:**
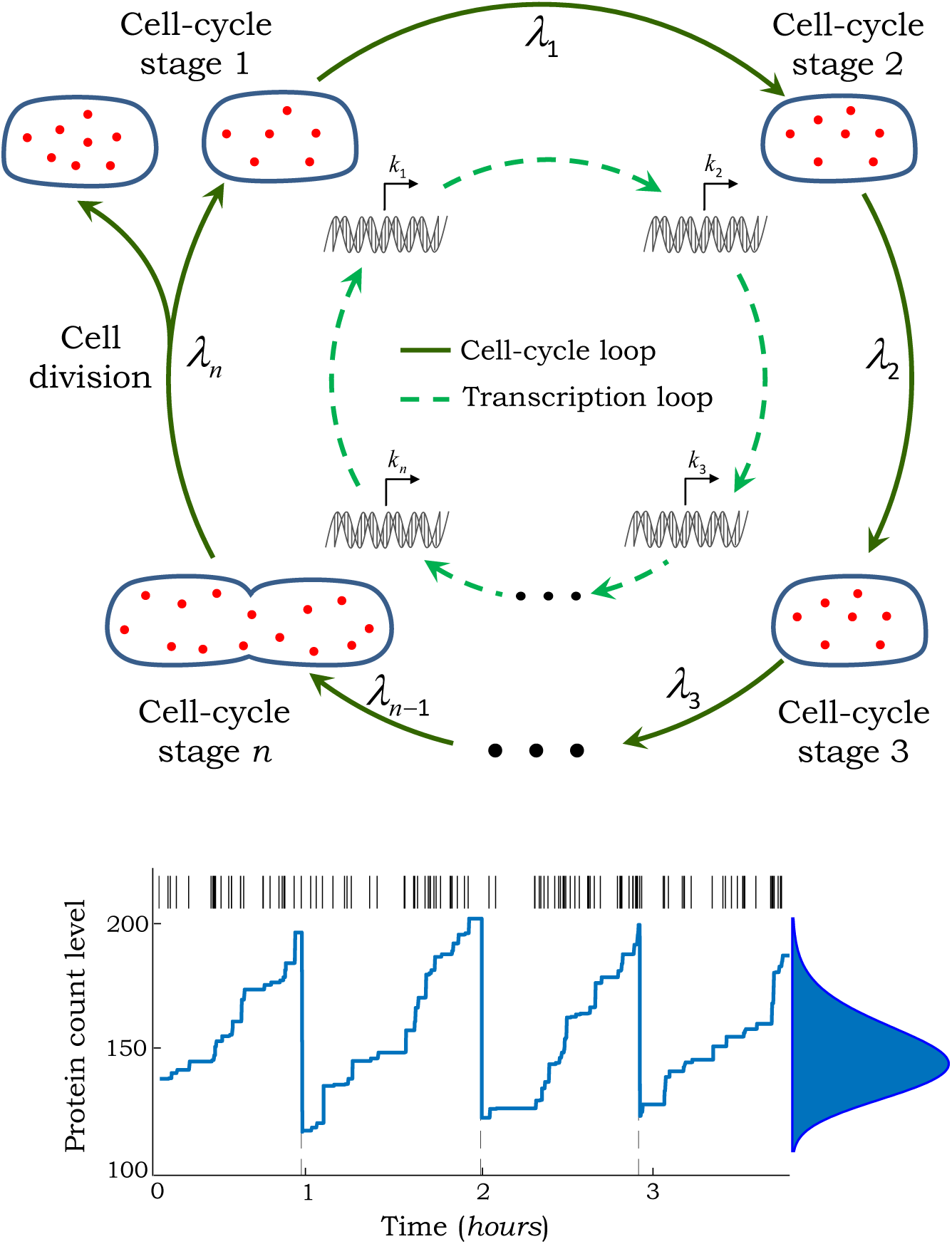
Coupling cell cycle to gene expression. *Top:* The outer loop shows an individual cell from birth to division passing through cell-cycle stages *C*_1_, *C*_2_,…, *C*_n_, with transition rates between stages give by λ_i_, *i* ∈ {1,2,…, *n*}. The cell is born in stage *C*_1_ and division is initiated in *C*_*n*_. The inner loop (transcriptional cycle) represents the rate at which protein expression bursts occur and is given by *k*_*i*_ in cell-cycle stage *C*_*i*_. *Bottom:* Representative trajectory of the protein level in an individual cell through multiple cell cycles (dashed lines). In this case, the transcription rate is assumed to double at the cell-cycle midpoint due to replication-associated increase in gene dosage. The spike train above represents the firing times of burst events. Steady-state distribution of the protein copy numbers obtained from running a large number of Monte Carlo simulations is shown on the right. The cell cycle was modeled by choosing *n* = 20 stages with equal transition rates between. Protein expression was assumed to occur in geometric bursts with *〈B〉* = 10 and molecules were partitioned between daughter cells based on a binomial distribution.

In the proposed model, some cell-to-cell variability or noise in the protein level is simply a result of cells being in different cell-cycle stages (i.e., asynchronous population). We illustrate a novel approach that takes into account such cell-cycle effects, and quantifies the noise contribution just from bursty expression and partitioning errors. Formulas obtained using this approach reveal that cell-cycle dependent expression considerably alters noise levels, always affecting contributions from bursty expression and partitioning errors in opposite ways. Intriguingly, our results show existence of optimal strategies to synthesize a protein within the cell cycle that minimize noise contributions for a fixed mean protein level. For example, the noise contribution from bursty expression is minimal when the protein is synthesized only towards the end of cell cycle. We discuss intuitive reasoning behind these optimal strategies, and provide examples of proteins that are expressed in this fashion to enhance fidelity of cell-cycle decisions.

## 2. Model coupling cell cycle to gene expression

We adopt a phenomenological approach to model cell cycle and divided it into *n* stages *C*_1_, *C*_2_, …, *C*_*n*_. A newborn cell is in stage *C*_1_ and transitions from *C*_i_ to *C*_*i+1*_ with rate λ_*i*_. In stage *C*_*n*_, cell division is initiated with rate λ_*n*_, and upon division the cell returns to *C*_1_. In the stochastic formulation of this model, the cell resides in stage *C*_*i*_ for an exponentially distributed time interval with mean 1/λ_i_, and cell-cycle duration is a sum of *n* independent, but not necessarily identical, exponential random variables. These stages can be mathematically characterized by Bernoulli processes *c*_1_(*t*), *c*_2_(*t*), …, *c*_*n*_(*t*), where *c*_*i*_(*t*) = 1 when the cell is in stage *C*_*i*_ and *c*_*i*_(*t*) = 0 otherwise. Based on the model structure, these processes satisfy

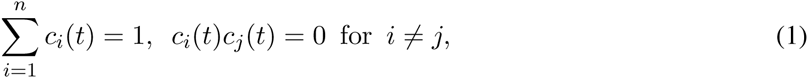

The latter equality results from the fact that only one of the *c*_*i*_ can be equal to 1 at any given time. In addition, since c_i_ takes values in {0,1}

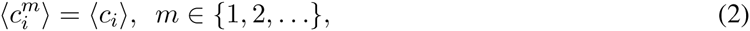

where the symbol 〈〉 denotes the expected value. Next, we describe the coupling between the cell cycle and stochastic expression models.

We assume that gene-expression bursts occur at aPoisson rate *k*_*i*_ in cell-cycle stage *C*_*i*_. Using the above-defined Bernoulli processes, the burst arrival rate can be compactly written as 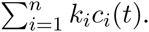 Let *x*(*t*) denote the number of protein molecules in a singe cell at time *t*. Then, whenever burst events occur, the protein level is reset as

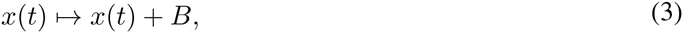

where the burst size *B* ∈ {0,1, 2,…} is a random variable independently drawn from an arbitrary distribution, and reflects the net contribution of transcriptional and translational bursting. As is true for most proteins in *E. coli* and *S. cerevisiae*, we assume a stable protein without any active degradation between burst events [43–45]. At the time of cell division (as dictated by the cell-cycle model), the protein molecules are randomly partitioned between daughters. This corresponds to the following reset that is activated during division

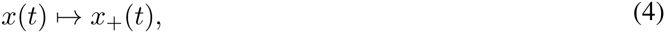

where the mean and variance of *x*_+_ (level just after division) conditioned on *x* (level just before division) are given by

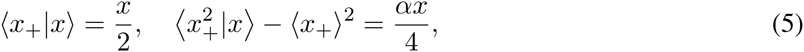

respectively. The first equation in (5) shows that the number of molecules is approximately halved during division, while the second equation quantifies the stochasticity in the partitioning process through the parameter *α*. The ideal case of zero partitioning errors corresponds at *α* = 0, where *x*_+_(*t*) = *x*(*t*)/2 with probability one. Binomial partitioning, where each molecule has an equal chance of ending up in one of the two daughter cells, is given by *α* = 1 [46–48]. Finally, values of *α* > 1 represent additional noise in the partitioning process that arise when protein molecules form multimers, or reside in organelles that are themselves subject to binomial partitioning [34,49]. The overall model coupling cell cycle to expression is illustrated in Fig. 1 together with a representative trajectory of *x*(*t*).

## 3. Mean protein level for cell-cycle driven expression

We illustrate an approach based on closing moment dynamics for deriving an exact analytical formula for the mean protein level. The first step is to obtain differential equations describing the time evolution of the statistical moments for *x*(*t*) and *c*_*i*_(*t*). These equations can be derived using the Chemical Master Equation (CME) corresponding to the stochastic model presented in the previous section (see Appendix A in SI). In particular, time evolution of the means (first-order moments) is given by

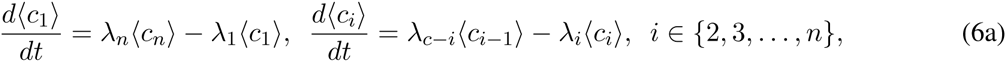

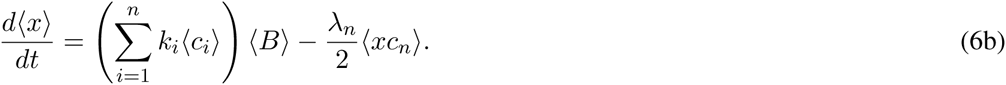

Steady-state analysis of (6a) yields the average value of Bernoulli processes as

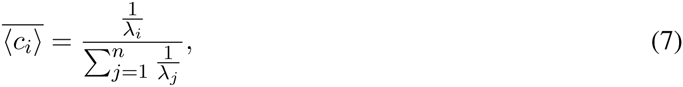

which can be interpreted as the fraction of time spent in the cell-cycle stage *C*_*i*_. We use 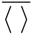 to denote the expected value of a stochastic process as *t* → ∞.

Note that the dynamics of 〈*x*〉 in (6b) is “not closed”, in the sense that it depends on second-order moments 〈*xc*_*n*_〉. This leads to the well-known problem of moment closure that often arises in stochastic chemical kinetics [50-57]. It turns out that in this case, the model structure can be exploited to automatically close moment equations. This is done by augmenting the system of equations in (6) with the time evolution of moments of the form 〈*xc*_*i*_〉

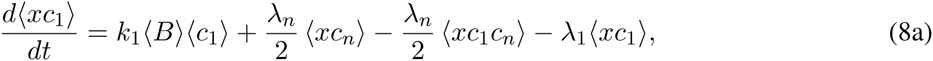

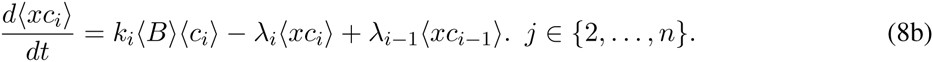

At the first look, these equations are unclosed and depend on third-order moments of the form 〈*xc*_*i*_*c*_*n*_〉. However, exploiting the fact that *c_i_c_j_ =* 0 from (1) leads to trivial closure

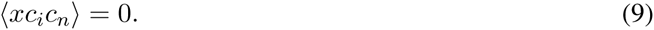

After using (9) in (8a), the mean protein level can be computed exactly by solving a linear dynamical system given by (6) and (8). At steady-state, the linear equations can be solved recursively to yield

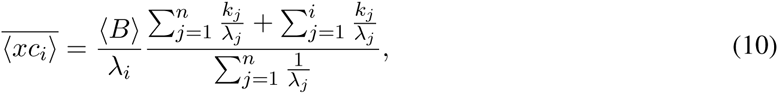

where 〈*B*〉 is the mean protein burst size. Since *c*_*i*_’s are binary random variables, the mean protein level conditioned on the cell-cycle stage (i.e., synchronized cell population) can be obtained as

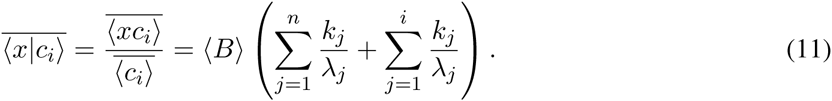

Furthermore, using (10) and the fact that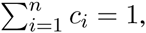

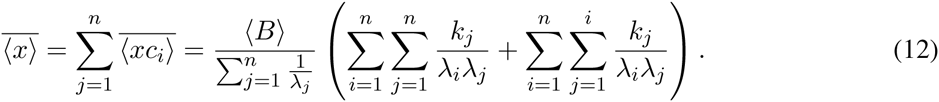

Next, we investigate the mean protein level 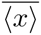 in some limiting cases. Consider equal transition rates between cell-cycle stages λ_*i*_ = *n*/*T*, which corresponds to an Erlang distributed cell-cycle durations with mean *T* and shape parameter *n*. In this scenario

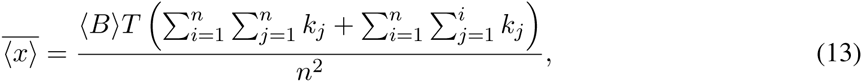

and further reduces to

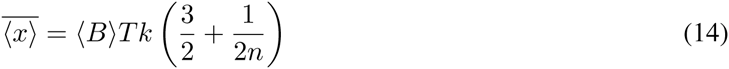

when the rate of expression bursts *k*_*i*_ = *k* is constant throughout the cell cycle. Finally, in the limit of deterministic cell-cycle durations of length *T*(*n* → ∞)

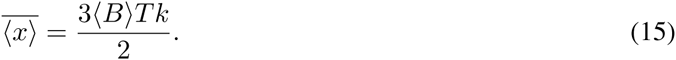

## 4. Protein noise level for cell-cycle driven expression

The mathematical approach illustrated above is now used to obtain the noise in protein copy numbers. By noise, we mean the magnitude of fluctuations in *x*(*t*) that can be attributed to two stochastic mechanisms: bursty expression and random partitioning. Note that even in the absence of these mechanisms, there will be cell-cycle related fluctuations with protein molecules accumulating over time and dividing by half at random cell-division times. To correct for such cell-cycle driven fluctuations, we define another stochastic process *y*(*t*) that estimates the protein level if expression and partitioning were modeled deterministically. More specifically, within the cell cycle *y*(*t*) evolves according to the following differential equation

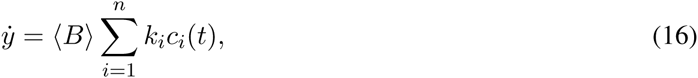

which is the deterministic counterpart to the stochastic expression model presented earlier. At the time of cell division, the level is divided exactly by half

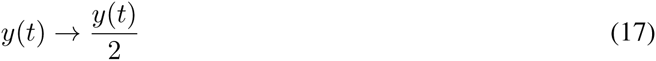

with zero partitioning errors, i.e., *α* = 0 in (5). This allows us to define a new zero-mean stochastic process *z*(*t*) corrected for cell-cycle effects

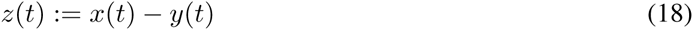

that measures the deviation in the protein count in the original stochastic model (*x*) from its expected levels if noise mechanisms were modeled deterministically (*y*). The protein noise level can now be defined through the dimensionless quantify

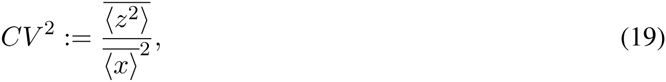

measuring the steady-state variance in *z*(*t*) normalized by the square of the mean level. Since 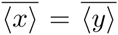 and 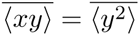 (see Appendix B in SI), it can be rewritten as

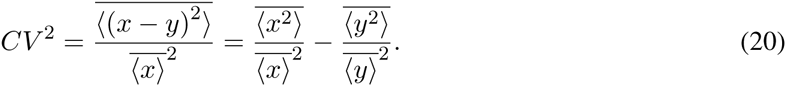

In the context of prior work, 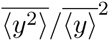 is interpreted as the extrinsic noise in gene expression resulting from cell-cycle effects. It is typically measured by the covariance in the singe-cell expression of two identical copies of a gene with common cell-cycle regulation [58,59]. In contrast, *CV*^2^ is the “intrinsic noise” resulting from stochasticity in gene expression and partitioning processes, and is measured by subtracting the extrinsic noise from the total noise 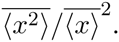

Having appropriately defined the noise level, we next compute it using moment equations. The time evolution of the moments 〈*z*^2^〉 and 〈*z*^2^*c*_*i*_〉 are given by (see Appendix C in SI)

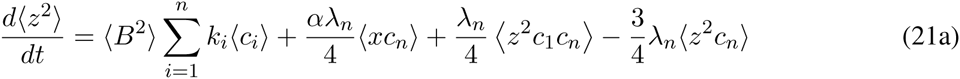

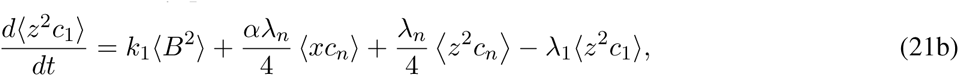

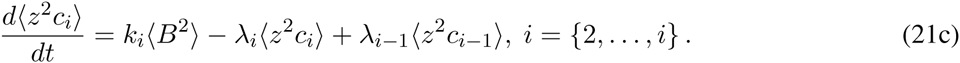

and depend on the fourth-order moments 〈*z*^2^*c*_1_*c*_*n*_.〉 Exploiting the model structure as before, it follows from (1) that 〈*z*^2^*c*_1_*c*_*n*_ = 0〉, and (6), (8), (21) constitute a “closed” set linear differential equations. Steady-state analysis yields the following noise level (see Appendix C in SI)

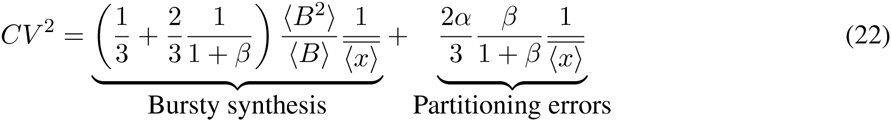

that is inversely proportional to the mean 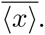 The noise can be decomposed into two terms: the first term represents the contribution from protein synthesis in random bursts and depends on the statistical moments of the burst size *B*. The second term is the contribution from partitioning errors and depends linearly on *α*. Recall that *α* measures the degree of randomness in partitioning of molecules between daughter cells, and is defined through (5). Interestingly, results show that the effect of cell-cycle regulation on the noise level can be quantified through a single dimensionless parameter

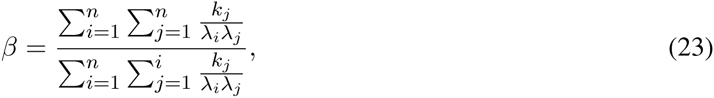

that is uniquely determined by the number of cell-cycle stages in the model (*n*), transition rates between stages (λ_*i*_), and protein synthesis rates across stages (*k*_*i*_). Note from (22) that *β* affects the noise terms in opposite ways-any coupling of cell-cycle to expression that increases *β* will attenuate the contribution from bursty expression but amplifies the contribution from partitioning errors. Finally, we point out that in the case of non-bursty expression (*B* = 1 with probability one) and binomial partitioning (*α* = 1)

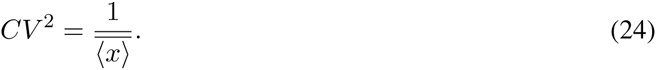

and the noise level is always consistent with that of a Poisson distribution^1^ irrespective of the value of *β*, and hence the form of cell-cycle regulation.

## 5. Optimal cell-cycle regulation to minimize noise

We explore how different forms of cell-cycle regulation affect *CV*^2^ and begin with the simplest case of a constant synthesis rate *k*_*i*_ = *k, i* ∈ {1, 2, …, *n*} throughout the cell cycle. This case would correspond to a scenario where the net rate of expression (across all copies of a gene) remains invariant to replication-associated changes in gene dosage, as has recently been shown in different organisms [32, 33]. Further assuming equal transition rates λ_*i*_ = *n*/*T* (Erlang distributed cell-cycle durations)

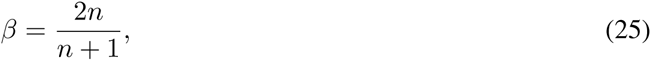

which reduces to *β* = 2 as *n* → ∞. Thus, in this important limit of no cell-cycle regulation (equal *k*_*i*_’s) and deterministic cell-cycle duration (large *n*),

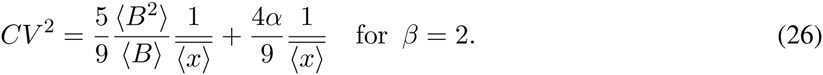

Next, consider the following strategies for coupling cell cycle to gene expression:

1. The burst arrival rate is assumed to increase by two-fold at the cell-cycle midpoint due to gene duplication. Assuming even *n*, this corresponds to

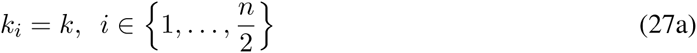

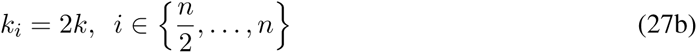
2. Expression only occurs at the start of cell cycle, i.e., *k*_1_ = *k* and all other *k*_*i*_’s are zero.
3. Expression only occurs at the end of cell cycle, i.e., *k*_*n*_ = *k* and all other *k*_*i*_’s are zero.
4. Expression only occurs at the cell cycle midpoint, i.e., 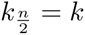 and all other *k*_*i*_’s are zero.

For a mathematically controlled comparison, the parameter *k* is adjusted using (12) from case-to-case so as to maintain a fixed average number protein molecules. The noise levels corresponding to the different forms of cell-cycle regulation are illustrated in Fig. 2. Interestingly, duplication of the protein expression rate within the cell cycle leads to a lower noise contribution from bursty synthesis, as compared to a constant rate throughout the cell-cycle. Moreover, expressing the protein only at the start (end) of cell cycle yields the highest (lowest) noise contribution from bursty synthesis. As expected from (22), the noise contribution from partitioning errors exhibits a completely opposite trend (Fig. 2).

**Figure 2:**
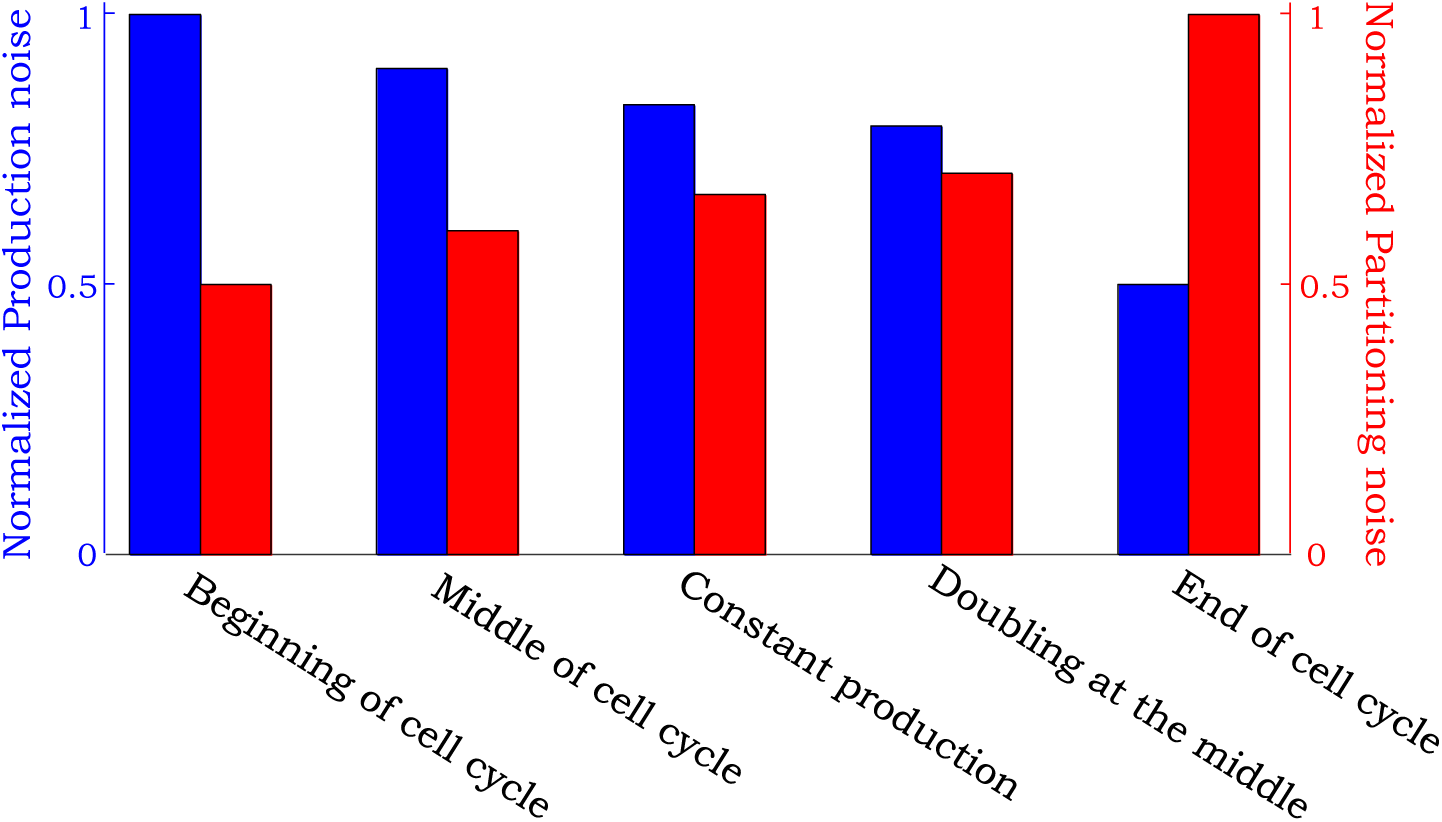
Noise comparison for different strategies coupling cell cycle to gene expression. The noise contributions from bursty expression (left) and partitioning errors (right) as given by (22) are shown for five different strategies: expression only at the start of cell cycle; expression only at the cell-cycle midpoint; constant mRNA synthesis rate throughout the cell cycle; doubling of synthesis rate at the cell-cycle midpoint; expression only towards the end of cell cycle. While noise contribution from bursty expression is minimized in the latter strategy, contribution from partitioning errors are lowest if expression occurs only at the beginning of cell cycle. The cell cycle was modeled by choosing *n* = 20 stages with equal transition rates and production rates *k*_*i*_ were chosen so as to have the same mean protein level per cell across all cases.

The above analysis begs an intriguing question: Is there an optimal way to express a protein during the cell cycle that maximizes/minimizes noise levels? Since the form of cell-cycle regulation impacts *CV*^2^ through *β*, this amounts to choosing *k*_*i*_’s so as to maximize/minimize it. Our result show that *β* is bounded from both below and above (see Appendix D in SI)

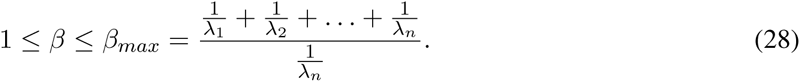

The minimal value of *β* = 1 is attained when expression only occurs at the start of cell cycle, i.e., a non-zero *k*_1_ and all other *k*_*i*_’s are zero. In this case

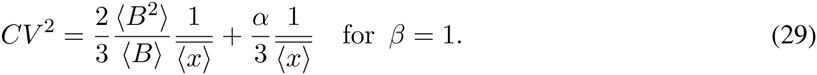

with the lowest noise contribution from partitioning errors, but the highest contribution from bursty synthesis. In contrast, the maximum value of *β* = *β*_*max*_ is attained when expression only occurs at the end of cell cycle, i.e., a non-zero *k*_*n*_ and all other *k*_*i*_’s are zero. Note form (28) that *β*_*max*_ → ∞ as *λ*_*n*_ → ∞ (time spent in stage *C*_*n*_ approaches zero), in which case

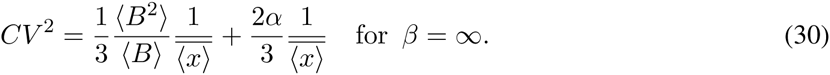

and the noise contribution from bursty synthesis is minimal.

In summary, if bursty expression is the dominant source of noise (high *B* and low *α*), then *CV*^2^ in minimized for a given 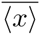 when the protein is made in the shortest time window just before cell division (Fig. 3). On the other hand, if randomness in partitioning error is dominant (low *B* and high *α*), the optimal strategy is to make the protein just after cell division. Finally, we point out that these optimal strategies also minimize stochastic variation in protein counts among synchronized cells, where all cells are in the same cell-cycle stage (see Appendix E in SI).

**Figure 3:**
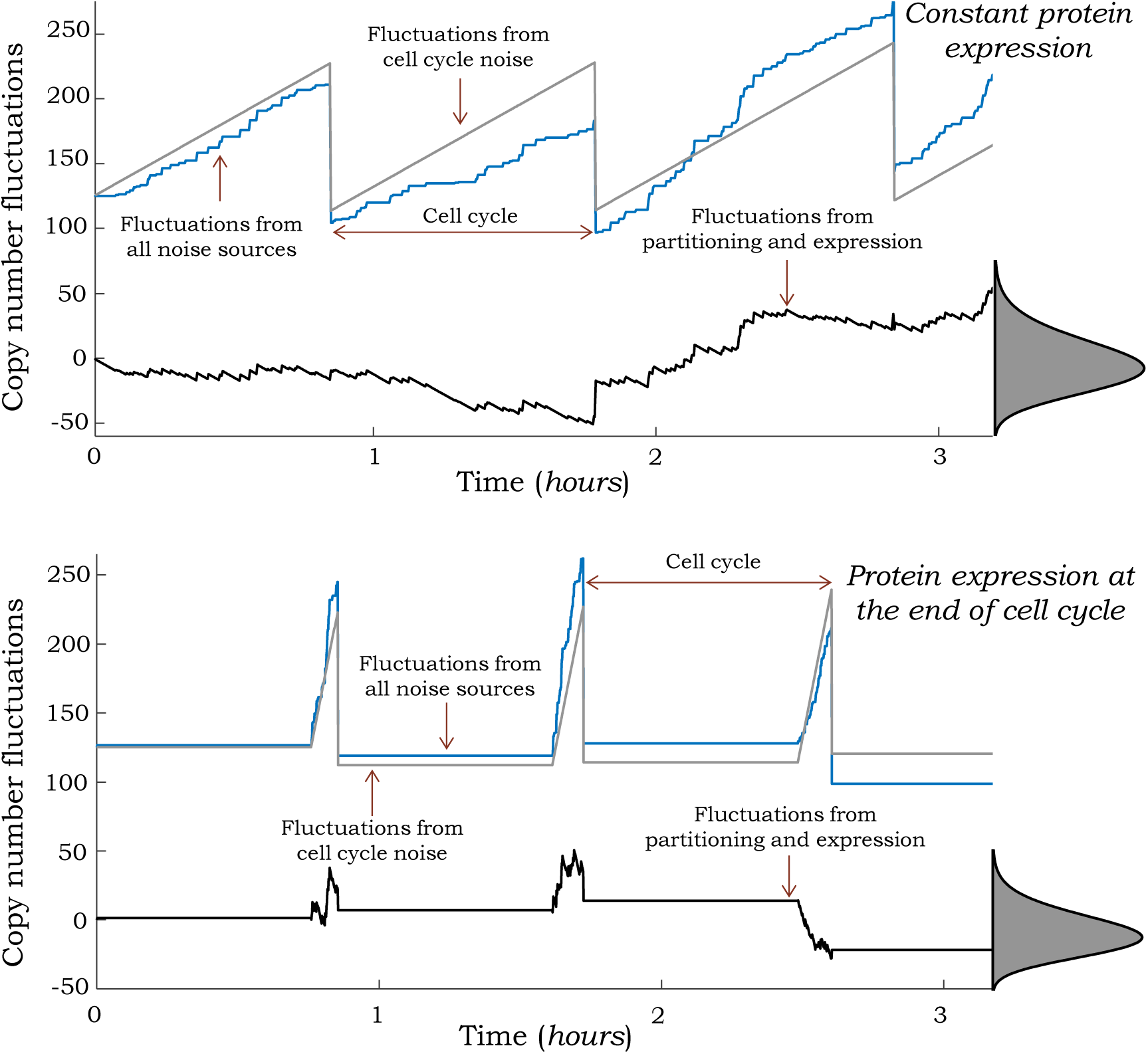
Synthesis of proteins towards the end of cell cycle minimizes fluctuations in copy numbers. Protein level in an individual cell across multiple cell cycles for two strategies: a fixed transcription rate throughout the cell cycle (top) and transcription only occurring just before cell division (bottom). Trajectories obtained via Monte Carlo simulations are shown for the stochastic model (blue) and a reduced model where noise mechanisms are modeled deterministically (gray). These levels are subtracted to obtain a zero-mean stochastic process *z*(*t*), where fluctuations resulting from cell cycle are removed (black). Steady-state distribution of *z* obtained from 10, 000 MC simulation runs is shown on the right, and the bottom strategy leads to lower variability in *z* for the same mean protein level. Cell cycle and expression was modeled as in Fig. 1 and burst arrival rates were chosen so as to ensure a average protein copy number of 150 molecules per cell in both cases.

## 6. Discussion

Theoretical model of stochastic gene expression have played a pivotal role in understanding how noise mechanisms and biologically relevant parameters generate differences in protein/mRNA population counts between isogenic cells [60-66]. Here we have expanded this theory to consider cell-cycle regulated genes. Our approach involves a general model of cell cycle, where a cell transitioning through an arbitrary number of stages from birth to division. The protein is assumed to be expressed in random bursts, and the rate at which bursts arrive varies arbitrarily with cell-cycle stage. In the case of translational bursting of proteins from mRNA, the burst arrive rate corresponds to the mRNA synthesis (transcription) rate. In contrast, for transcriptional bursting of mRNAs, the burst arrive rate corresponds to the frequency with which a promoter become transcriptionally active. The key contribution of this work is derivation of (12) and (22) that predict the protein mean and noise levels for a given form of cell-cycle regulation.

Derivation of noise formulas enable uncovering of optimal cell-cycle regulation strategies to minimize *CV*^2^ for a fixed mean protein level. In the physiological case of large bursts (*B* ≫ 1) and binomial partitioning of proteins between daughter cells (*α* = 1), the contribution from bursty synthesis dominates *CV*^2^. Our results show that in this scenario, expression of the protein just before division is the optimal strategy (Fig. 3). Intuitively, such a strategy can be understood in the context of the number of burst events from birth to division needed to maintain a given 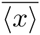 throughout the cell-cycle. It turns out that this number is highly dependent on the form of cell-cycle regulation. Hence, any strategy that requires more burst events to maintain the same mean protein level, lowers noise through more effective averaging of the underlying bursty process, albeit being more energy inefficient. For example, if protein production only occurs at the end of cell cycle, then on average, 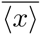 number of proteins have to added just before cell division. This corresponds to 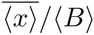 number of burst events per cell cycle. If proteins were only expressed at the start of cell cycle, then one needs to add only 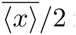 number of molecules, half as much as the earlier strategy. If proteins were made at a constant synthesis rate throughout the cell cycle, then on average, 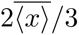 number of protein are added per cell cycle, which is higher than the early-expression strategy but lower than the late-expression strategy. In summary, gene product synthesis just before division requires production of the most number of protein molecules to maintain a fixed mean level within the cell-cycle, and hence, provides the most effective noise buffering through averaging of burst events. Next, we provide two recent examples of proteins that are indeed expressed in this fashion.

The green alga *C. reinhardtii* has a prolonged *G*_1_ phase, where the size of a newborn cell increases by more than 2-fold. This long *G*_1_ phase is followed by an *S/M* phase. Here the cell undergoes multiple DNA replication and fission cycles creating *2^d^* daughter cells, where *d* is number of rounds of division. Recent studies suggest that the number of rounds of division is controlled by a protein CDKG1, that is only expressed just before exit from *G*_1_ [67]. Another example, is the protein Whi5 in budding yeast *S. cerevisiae* and its level controls the transition of cells past the Start checkpoint. This protein in not expressed in *G*_1_, and is only synthesizes late in the cell cycle [68,69]. While such selective expression of these proteins plays a critical role in coupling cell size to cell-cycle decision, it may also minimize intrinsic fluctuations in protein levels from the innate stochasticity in gene expression. Clearly, a more systematic study exploring the role of noisy expression on the fidelity of these cell-cyle decisions is warranted.

It is important to point out that our analysis made various simplifying assumptions, such as, i) Excluding time evolution of cell size and size-dependent expression; ii) Instantaneous transcriptional and translational bursts that correspond to short-lived mRNAs and active promoter states; iii) Cell-cycle durations being independent random variables, implying no correlation between the division times of mother and daughter cells. While many of these assumptions are clearly violated for cellular systems, they were necessary to obtain exact analytical solutions that provide novel insights into noise control by synchronizing gene expression to cell cycle. Further work will focus on relaxing these assumptions, in particular, the first assumption of incorporating cell size into the model. This will allow investigation of both concentration and copy number of gene products in single cells, and some recent work on modeling stochastic dynamics of cell size has already been done [70-72].

## Author contributions

AS defined the problem and formulated the approach. MS did the mathematical derivations and both authors collaborated on writing the paper.

## Acknowledgments

AS is supported by the National Science Foundation Grant DMS-1312926. We thank Cesar Vargas-Garcia and Khem Ghusinga for feedback on the manuscript.

## Appendix A Moment equations describing the model

Based on standard stochastic formulation of chemical kinetics [73,74], the model describing *x* contains the following stochastic events

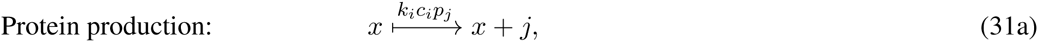

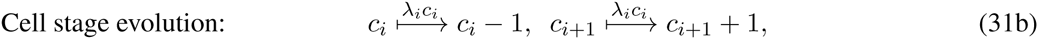

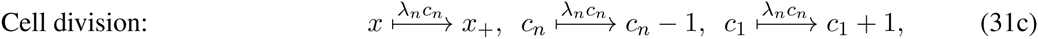

where the probability of having a burst of *j* molecules is given by *p*_*j*_. Whenever an event occurs, the states of the system change based on the stochiometries given in (31). On top of the arrows we showed the event propensity function *ψ*(*x,c*_*i*_), which determines how often reactions occur, i.e., the probability that an event occurs in the next infinitesimal time interval (*t*, *t* + *dt*] is *ψ*(*x*, *c*_*i*_)*dt*. Time derivative of the expected value of any function *φ*(*x*, *c*_*i*_) for this system can be written as [75]

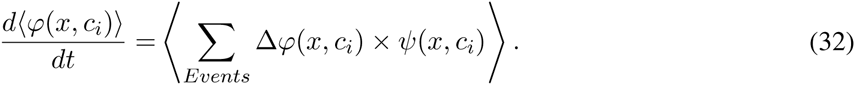

Choosing φ to be *x* and *c*_*i*_, *i* = {1,2,…, *n*} results in the equation (6) in the main article.

## B Moment dynamics of *y*

The model describing *x* and *y* includes the stochastic events

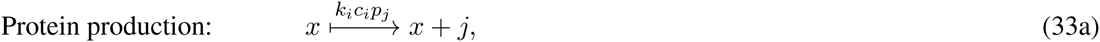

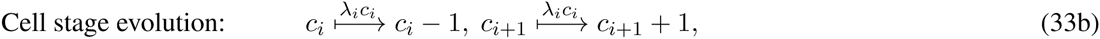

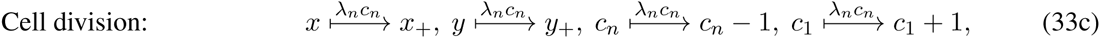

and the deterministic production of *y*

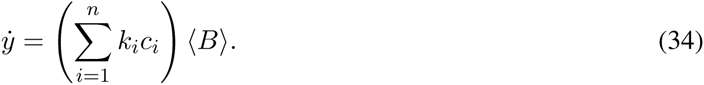

Time derivative of the expected value of any function *φ*(*x, y, c*_*i*_) for this system can be written as [75]

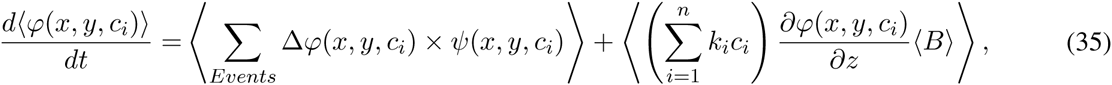

where the first term in the right-hand side is contributed from stochastic events and the second one is contributed from (34). The propensity function of events is given by *ψ*(*x, y, c*_*i*_). The mean dynamics of *y* can be written by choosing φ to be *y*

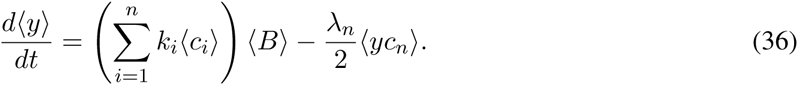

Dynamics of 〈*y*〉 is not closed and depends to moments 〈*yc*_*n*_〉, hence in order to have a closed set of equations we add new moments dynamics by selecting φ to be *yc*_*i*_

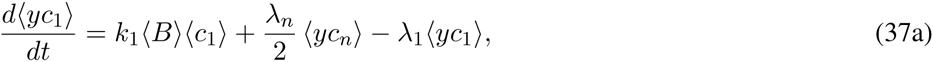

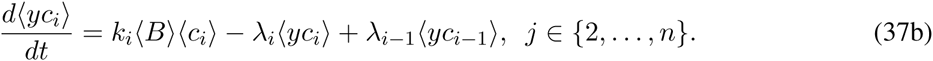

Dynamics of 〈*y*〉 and 〈*yc*_*i*_〉, *j* ∈ {1,…, *n*} are the same as dynamics of 〈*x*〉 and 〈*xc*_*i*_〉, *j* ∈ {1,…, *n*} presented in (6b) and (8) in the main text, hence 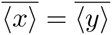 and 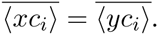.

Further, dynamics of 〈*xy*〉 can be written as

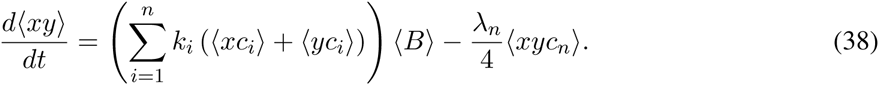

In order to have a closed set of equations we add dynamics of 〈*xyc*_*i*_〉

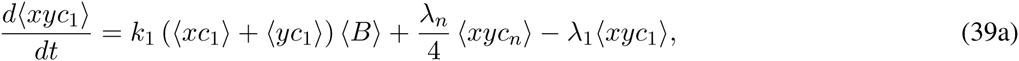

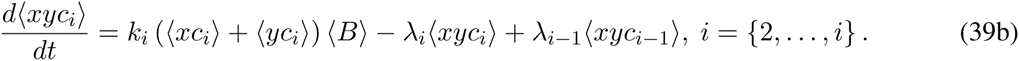

By having a closed set of equations related to *xy*, in the next step we add dynamics of 〈*y*^2^〉 and 〈*y*^2^*c*_*i*_〉

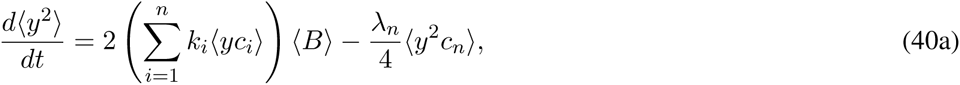

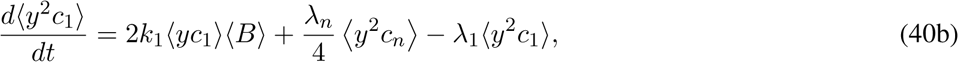

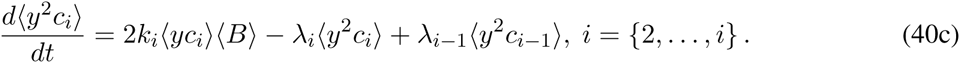

Using the fact that 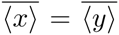 and 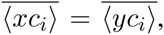 equations (40), (38), and (39) in steady-state results in 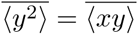 and 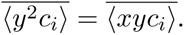

## C Calculation of z^2^

The random variable z is governed via

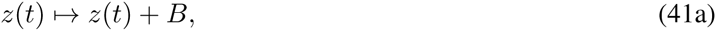

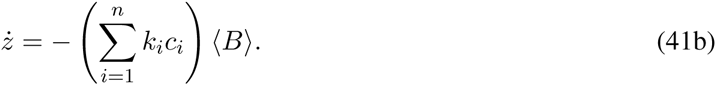

Further in the time of division, *z*_+_ is defined as

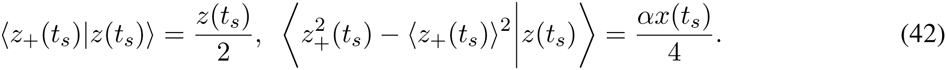

Hence the model by taking into account *z* contains the following stochastic events

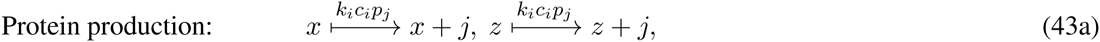

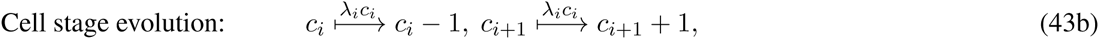

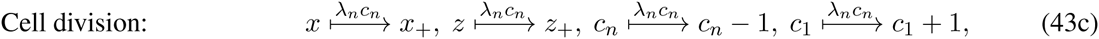

and deterministic dynamics of *z* given in (41b). Time derivative of the expected value of any function *φ*(*x, z, c*_*i*_) for this system can be written as [75]

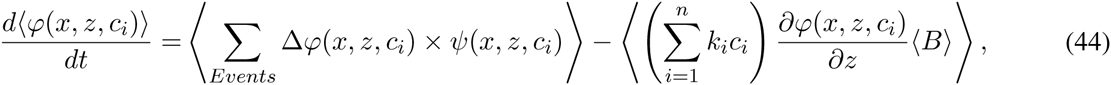

where the first term in the right-hand side is contributed from stochastic events and the second one is contributed from (41b). The propensity function of events is given by *ψ*(*x, z, c*_*i*_).

By choosing *φ* to be *z*^2^ and *z*^2^*c*_*i*_, *i* = {1,…, *i*) we have the following moment dynamics

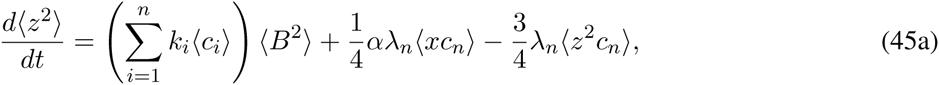

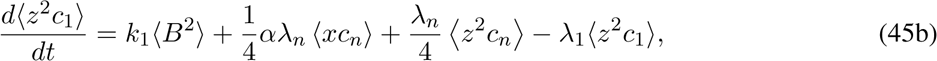

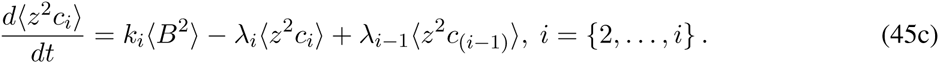

Note that just one of the binary states *c*_*i*_ can be 1 at a time, thus 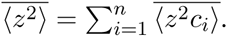. In order to calculate the terms 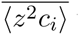 we need to express the term 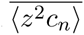 as the first step. This term can be calculated by analyzing equation (45a) in steady-state

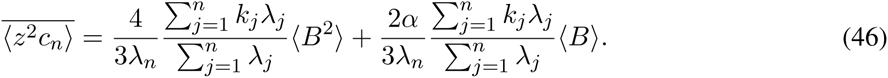

By using a recursive process we calculate moments 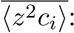 we calculate 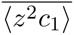 by substituting equation (46) in equation (45b). Then we use the definition of 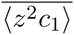 to calculate 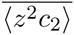 from equation (45c) and so on

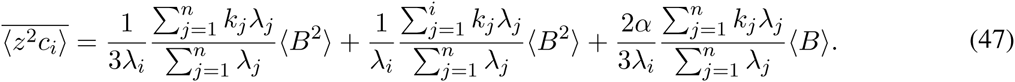

Summing up all the term in equation (47) results in 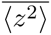

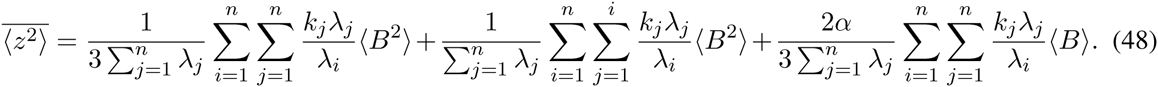

Finally, protein noise level can be written as

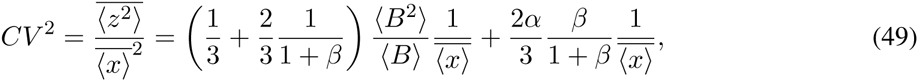

where

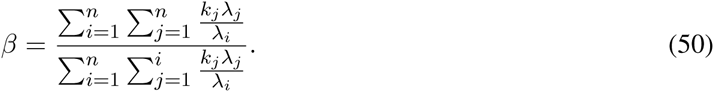

## D Optimal value of *β*

From (49) it is clear that minimum production noise occurs when *β* is maximum, and minimum value of partitioning noise happens when *β* is minimum. *β* can be written as

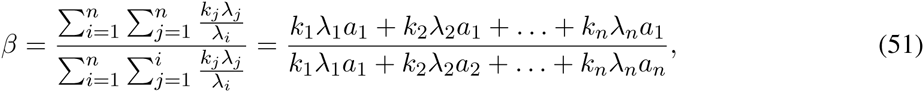

where

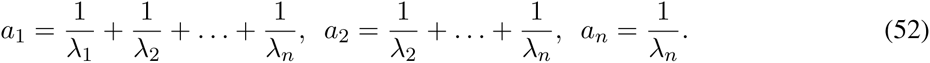

Note that

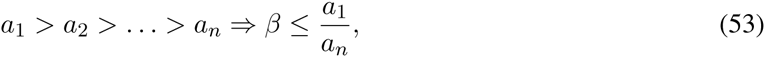

where equality happens when all *k*_*i*_s are zero except *k*_*n*_. Using the same methodology one can see that minimum of *β* happens when all the rates are zero except *k*_1_. The minimum value of *β* is one.

## E Noise in synchronized cells

Statistical moments conditioned on the cell cycle stage *c*_*i*_ can be obtained using

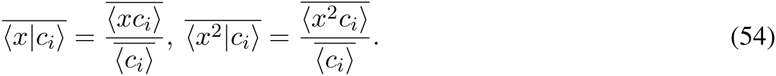

In order to calculate noise in synchronized cells we need to calculate 〈*x*^2^*c*_*i*_〉

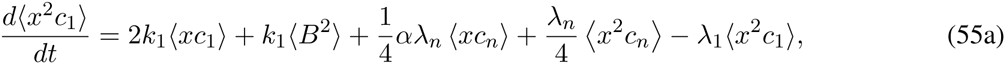

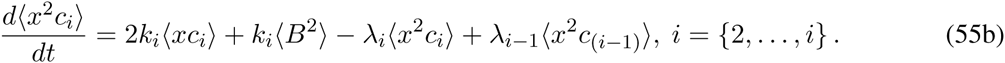

In order to calculate 〈*x*^2^*c*_*n*_〉 we introduce the moment dynamics of 〈*x*^2^〉

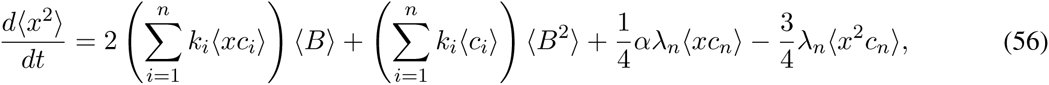

hence in steady-state

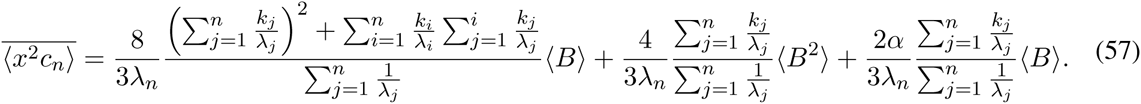

By using a similar process used in the previous section we calculate moments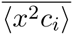

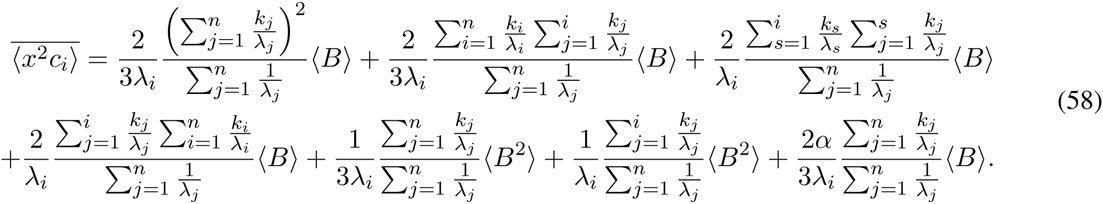

By having 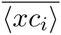 and 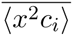 from (10) and (58), we can calculate mean and noise in synchronized cells. Using (54) yields the following conditional mean

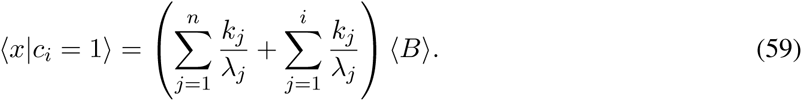

Further, the protein noise level given that cells are in stage *c*_*i*_ is given by

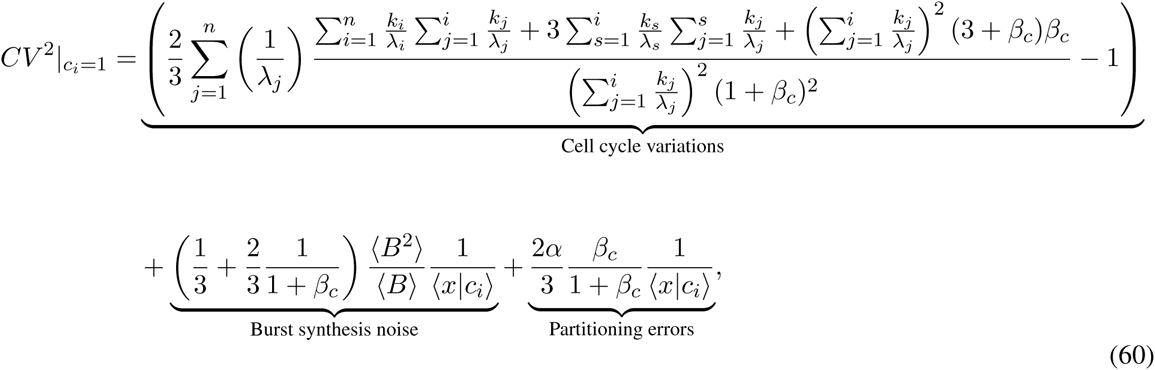

where

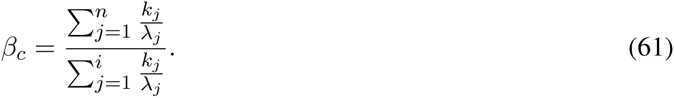

## Footnotes

1 The coefficient of variation squared for a Poisson distributed random variable is inverse of its mean

